## Supplementary information for "Structural insights into the elevator-type transport mechanism of a bacterial ZIP metal transporter"

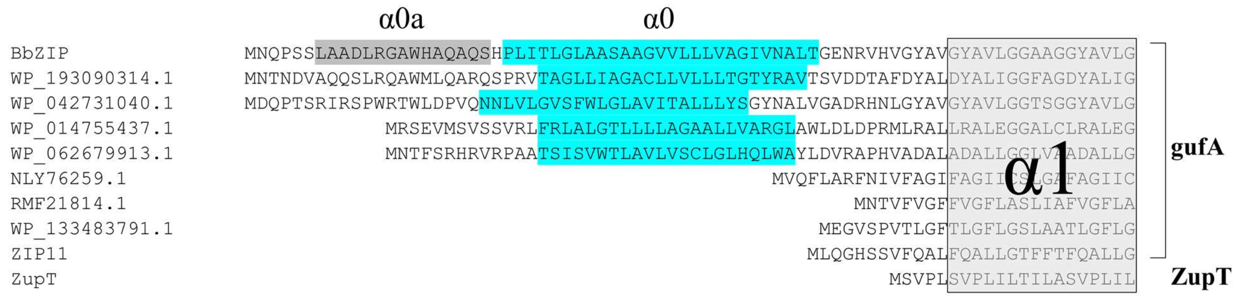


Figure S1. **Sequence alignment of the N terminal sequences of selected ZIPs.** BbZIP, RMF21814.1 [*Cyanobacteria bacterium J083*], NLY76259.1 [*Firmicutes bacterium*], WP_133483791.1 [*Halomonas ventosae*], WP_062679913.1 [*Achromobacter denitrificans*], WP_014755437.1 [*Pseudomonas*], WP_193090314.1 [*Advenella sp. FME57*], WP_042731040.1 [*Pseudomonas*], and human ZIP11 (Q8N1S5.3) are within the gufA subfamily. ZupT from *Escherichia coli* is a representative member of the ZupT subfamily. The amphipathic helix (α0a) and α0 of BbZIP are highlighted in grey and cyan, respectively. The predicted α0 of the other ZIPs, if there is any, are also highlighted. Note that some ZIPs in the gufA subfamily don't have α0 or α0a.

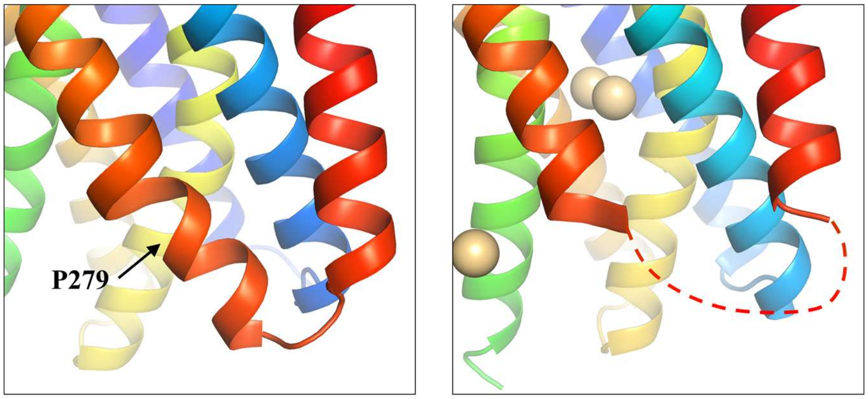


Apo state

$Cd^{2+}$ bound state
(5TSB)

Figure S2. **Structure of the loop connecting α7 and α8.** In the apo state structure (*left*), the residues missed in the previous structure (*right*) form a broken helix and a short loop. The conserved proline residue (P279) causes a kink in α7.

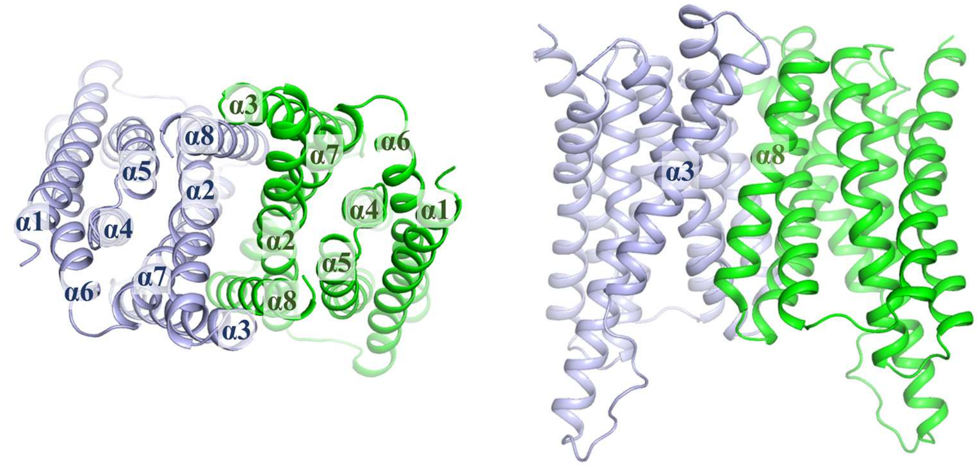


Figure S3. **BbZIP dimer predicted by AlphaFold in top view (*left*) and side view (*right*).**

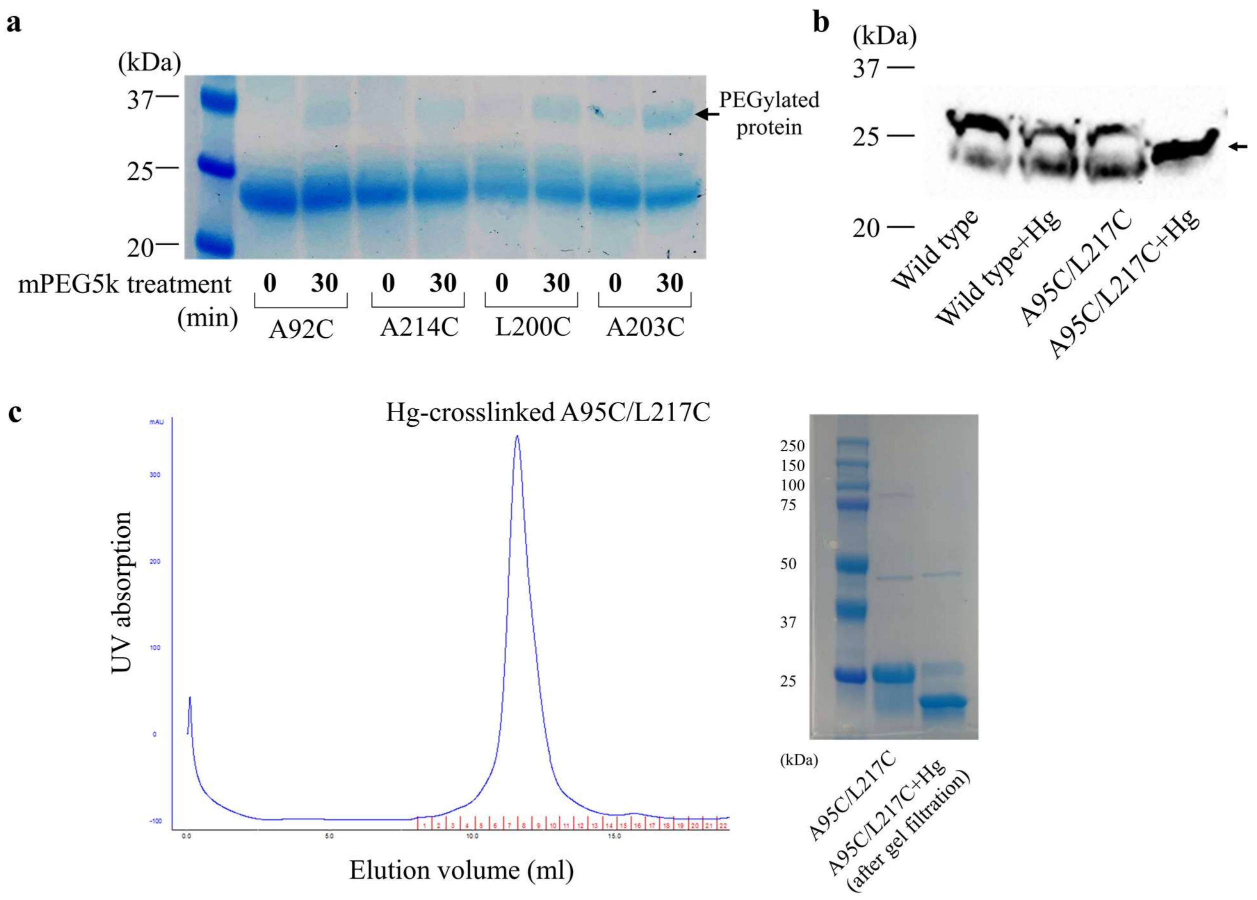


Figure S4. **Additional evidence supporting the proposed OFC model. a** Cysteine accessibility assay. Selected single cysteine variants were purified in DDM and treated with mPEG5K. EDTA was added to the sample immediately before the treatment to prevent cysteine blockage by $Cd^{2+}$. The reaction was terminated by 100 mM water soluble thiol reacting reagent methyl methanethiosulfonate before analysis in SDS-PAGE. The upper band indicated by the arrow is the PEGylated protein. **b** Hg-mediated chemical crosslinking of the A95C/L217C variant in the native membrane. The membrane fraction of the cells expression the variant was incubated with $HgCl_2$, terminated by NEM, and applied to Western blot by using a customized monoclonal antibody against BbZIP generated by Creative Biolabs Inc. The arrow indicates the crosslinked product. **c** Size-exclusion chromatography of the Hg-crosslinked of A95C/L217C variant. The eluted sample was applied to SDS-PAGE, and the band shift indicates that the protein is still in the crosslinked state after removal of free $Hg^{2+}$ from the sample.

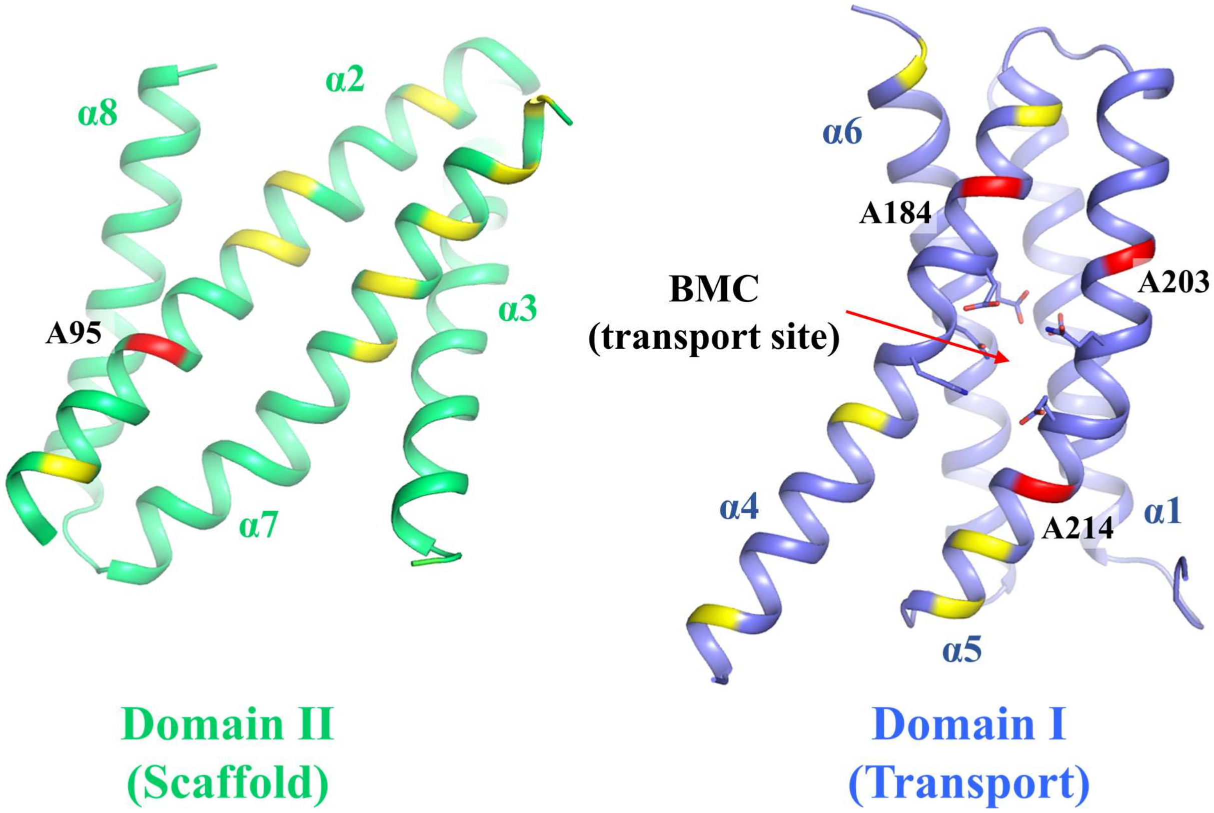


Figure S5. **Mapping of small residues at the interface between the transport domain and the scaffold domain.** The small residues (Gly, Ala, Ser) at the domain interface are colored in red (highly conserved) or yellow (less or non-conserved). The residues in the BMC (transport site) are shown in stick mode.

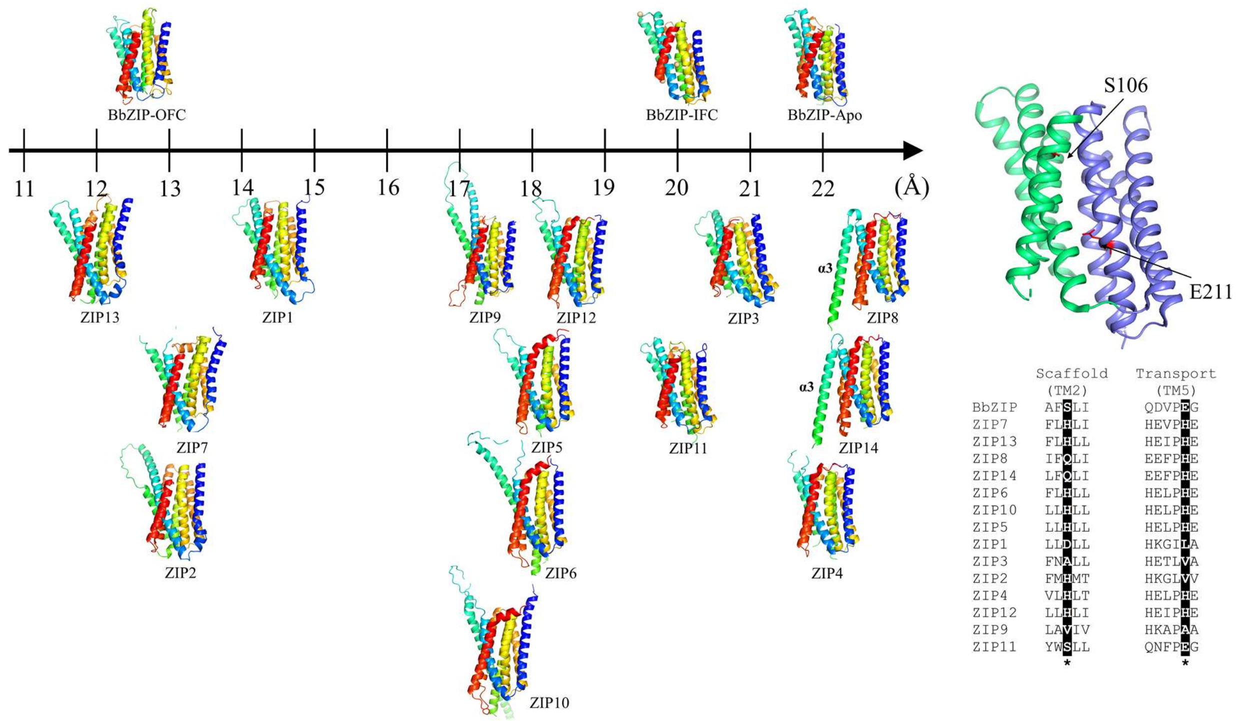


Figure S6. **Comparison of the BbZIP structures with the structures of human ZIPs predicted by AlphaFold**. The scaffold domains (α2/3/7/8) of the ZIPs are structurally aligned. For clarity, the extracellular domains and the cytosolic loop between α3 and α4 are not shown. To better distinguish conformational states, the structures are plotted against the distance between a residue at the pore entrance (S106 in BbZIP) and the last residue of the metal chelating motif in α5 (E211 in BbZIP). S106 and E211 are labeled in the BbZIP structure in the apo state (right upper corner) with the scaffold domain colored in green and transport domain in blue. The residue pairs in other ZIPs for distance measurement are highlighted in the sequence alignment and indicated with asterisks. Note that α3 of ZIP8 and ZIP14 is predicted to be adjacent to α8, which is the position for α3 of the other protomer when the transporter forms a homodimer as shown in Figure 5B and Figure S3.

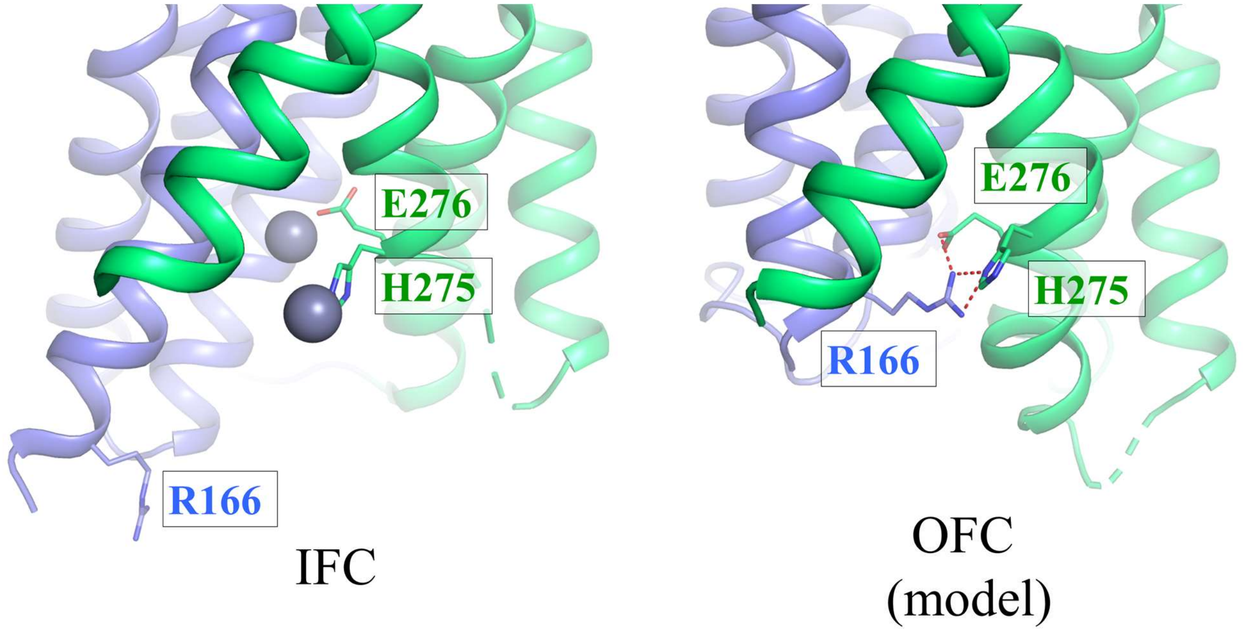


Figure S7. **A putative cytoplasmic gate in the OFC.** *Left*: IFC (PDB: 5TSA). *Right*: the OFC model. The residues involved in gate formation are labeled and shown in stick mode and. Zinc ions in the IFC are depicted as grey spheres. The hydrogen bonds are shown as red dashed lines. The transport domain and the scaffold domain are colored in blue and green, respectively.

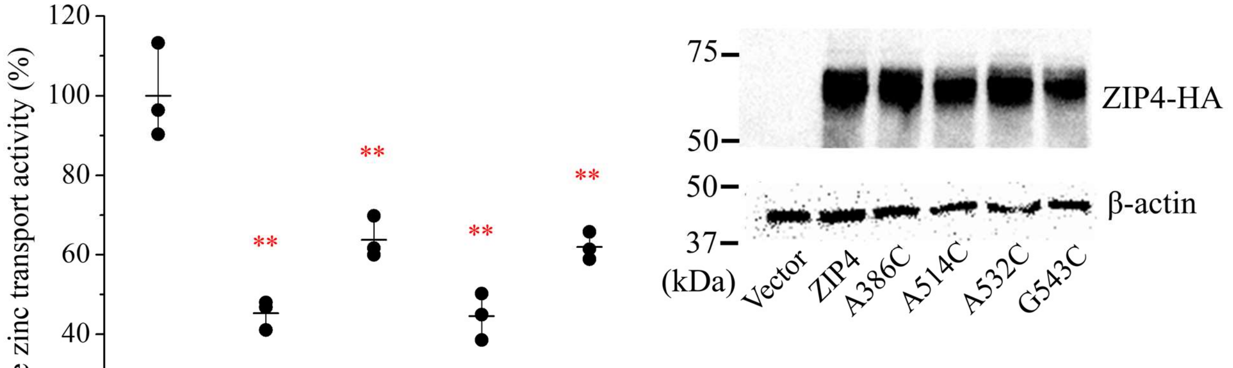


Normalized expression level

| ZIP4 | A368C | A514C | A532C | G543C |
|---|---|---|---|---|
| 1.0 | 1.2 | 1.2 | 1.4 | 1.1 |

Figure S8. **Functional analysis of the ZIP4 variants with cysteine substitutions.** A portion of cysteine substitutions in cysteine accessibility assay and crosslinking experiments was conducted on conserved residues of BbZIP, including A95, A184, A203, and A214. The corresponding residues in human ZIP4 were replaced with cysteine and the result variants were subjected to zinc transport assay. The relative transport activity of each variant is expressed as the percentage of the activity of the wild type ZIP4. The activity has been calibrated using the expression level estimated by Western blot, which is expressed as the normalized ratio of band intensity of HA-tagged ZIP4 to β-actin. The shown data are from one representative experiment and 2-3 independent experiments were conducted for each variant. Three replicates were included in one experiment. The horizontal bar of the scatter dot plot represents the mean and the vertical bar indicates the standard deviation. The asterisks indicate the significant differences between the variants and the wild type ZIP4 (Student's *t*-tests: ** $P \leqslant 0.01$).

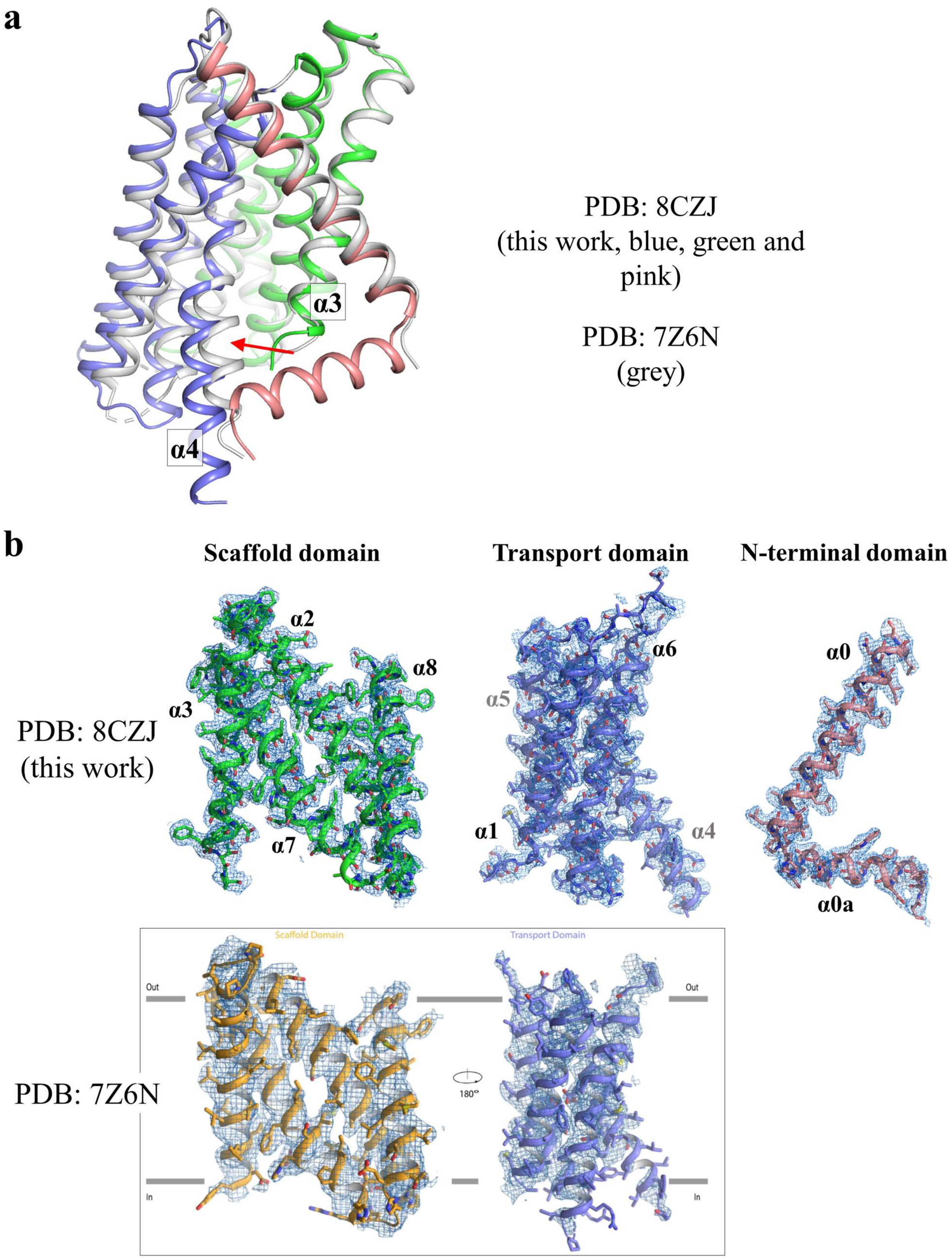
a
α3
α4
PDB: 8CZJ
(this work, blue, green and pink)
PDB: 7Z6N
(grey)
b
Scaffold domain
Transport domain
N-terminal domain
PDB: 8CZJ
(this work)
α2
α8
α3
α7
α6
α5
α1
α4
α0
α0a
PDB: 7Z6N
Scaffold Domain
Transport Domain
Out
In
180°

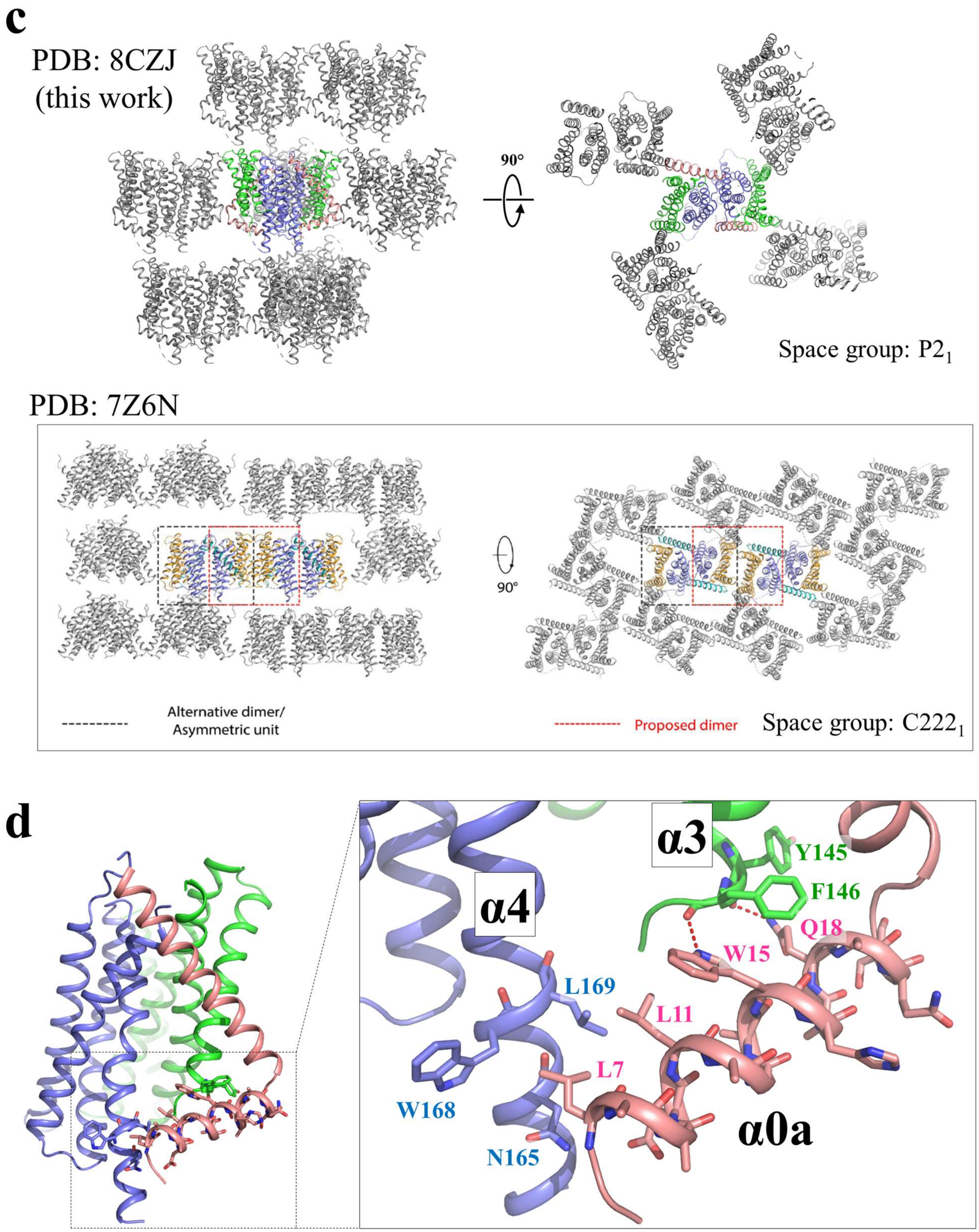


Figure S9. **Comparison of the structure solved in this work (PDB entry ID: 8CZJ) and the reported BbZIP structure in metal free state (PDB entry ID: 7Z6N). a** Structural comparison after alignment of the scaffold domains. The scaffold domain, the transport domain, and the N-terminal domain of 8CZJ are colored in green, blue and pink, respectively. 7Z6N (chain B) is shown in grey. Note that the orientation of the transport domains relative to the scaffold domain in the two structures are different with the cytoplasmic side of α4 being the most variable region as indicated by the arrow. **b** 2FoFc electron density maps (σ=1) of 8CZJ (2.75 Å) and 7Z6N (2.6 Å). The corresponding domains are structurally aligned with labeled structural elements. The density map of the N-terminal domain of 8CZJ, including the transmembrane helix α0 and amphipathic helix α0a, is also shown. The image of 7Z6N is adapted from Fig. S1B of Ref 82. **c** Comparison of crystal packing of 8CZJ with 7Z6N. The crystallographic dimer in one asymmetry unit of 8CZJ is colored and the symmetry mates are in grey. The image of 7Z6N in frame is adapted from Fig. S4 of Ref 82. **d** Association of α0a with α3 and α4 through hydrogen bonds (red dashed lines) and hydrophobic interactions, respectively. Hypothetically, α0a may limit the elevator-like movement of the transport domain (blue) relative to the scaffold domain (green) and therefore function as a negative regulator.

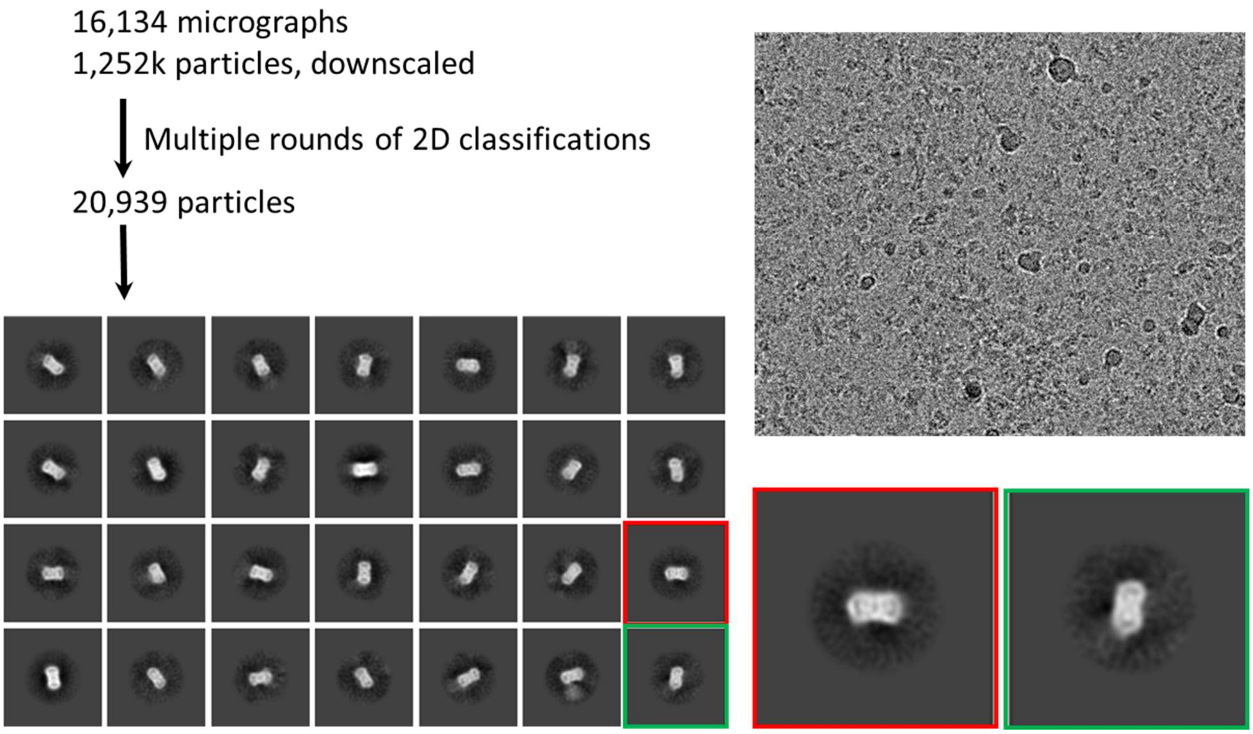


Figure S10. **The cryo-EM data processing workflow for BbZIP in Amphipol.**

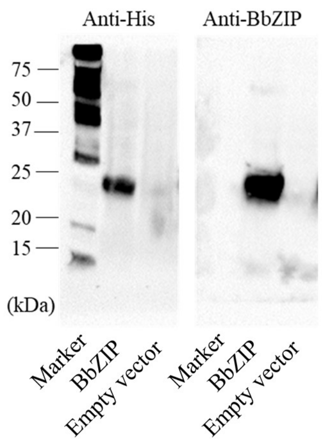


Figure S11. **Sensitivity and specificity of the customized monoclonal antibody against BbZIP.** The whole cell lysates of the cells transformed with the empty vector or the plasmid containing $His_6$-tagged BbZIP were applied to Western blot. The $His_6$-tagged BbZIP was detected with either an anti-His antibody (0.1 μg/ml) or the anti-BbZIP antibody (0.06 μg/ml). The result shows a better sensitivity and specificity of the anti-BbZIP antibody than the anti-His antibody.

**Table S1.** Crystallographic statistics

| **Crystal** | Apo BbZIP |
|---|---|
| **Data collection** | |
| Beamline | GM/CA-CAT (23-ID-D) |
| Wavelength (Å) | 1.0331 |
| Space group | $P\ 2_1$ |
| Unit cell | |
| a, b, c (Å) | 63.0, 117.7, 64.3 |
| α, β, γ (°) | 90, 104.2, 90 |
| [a]Resolution (Å) | 37.1 - 2.75 (2.85 - 2.75) |
| [a]Redundancy | 7.4 (2.6) |
| [a]Completeness (%) | 94.5 (74.8) |
| [a]$I/\sigma I$ | 7.8 (0.6) |
| [a,b]$R_{merge}$ | 0.187 (0.968) |
| [a,c]$R_{pim}$ | 0.068 (0.57) |
| [d]$CC_{1/2}$ of the highest resolution shell | 0.491 |
| **Refinement** | |
| Unique reflections | 22227 |
| Number of Atoms | 4289 |
| Protein | 4019 |
| Ligands | 263 |
| $H_2O$ | 7 |
| [e]$R_{work}/R_{free}$ | 0.233/0.261 |
| *Wilson B-factor* (Å$^2$) | 63.1 |
| *B*-factors (Å$^2$) | |
| Protein | 54.8 |
| MPG | 55.5 |
| $SO_4^{2-}$ | 78.2 |
| $H_2O$ | 52.3 |
| R.m.s. deviations | |
| Bond lengths (Å) | 0.010 |
| Bond angles (°) | 1.23 |
| Ramachandran plot (%) | |
| Favored | 97.3 |
| Allowed | 2.7 |
| Outliers | 0.0 |

[a]Highest resolution shell is shown in parentheses.

[b]$R_{merge} = \sum_{hkl} \sum_j |I_j(hkl) - \langle I(hkl) \rangle| / \sum_{hkl} \sum_j I_j(hkl)$, where $I$ is the intensity of reflection.

[c]$R_{pim} = \sum_{hkl} [1/(N-1)]^{1/2} \sum_j |I_j(hkl) - \langle I(hkl) \rangle| / \sum_{hkl} \sum_j I_j(hkl)$, where N is the redundancy of the dataset.

[d]$CC_{1/2}$ is the correlation coefficient of the half datasets.

[e]$R_{work} = \sum_{hkl} | |F_{obs}| - |F_{calc}| | / \sum_{hkl} |F_{obs}|$, where $F_{obs}$ and $F_{calc}$ is the observed and the calculated structure factor, respectively. $R_{free}$ is the cross-validation R factor for the test set of reflections (5% of the total) omitted in model refinement.

**Table S2.** Primers for mutagenesis in this work.

| Primers | Sequences (5'-3') |
|---|---|
| **BbZIP primers** | |
| H55C-forward | CTG ACC GGT GAA AAT CGT GTT TGC GTT GGT TAT GCA GTT CTG GGT |
| H55C-reverse | ACC CAG AAC TGC ATA ACC AAC GCA AAC ACG ATT TTC ACC GGT CAG |
| L92C-forward | GCA CGT ACC CAG GAT GCA ATG TGC GGT TTT GCC GCA GGT ATG ATG |
| L92C-reverse | CAT CAT ACC TGC GGC AAA ACC GCA CAT TGC ATC CTG GGT ACG TGC |
| A95C-forward | CAG GAT GCA ATG CTG GGT TTT TGC GCA GGT ATG ATG CTG GCA GCC |
| A95C-reverse | GGC TGC CAG CAT CAT ACC TGC GCA AAA ACC CAG CAT TGC ATC CTG |
| A102C-forward | GCC GCA GGT ATG ATG CTG GCA TGC AGT GCA TTT AGC CTG ATT CTG |
| A102C-reverse | CAG AAT CAG GCT AAA TGC ACT GCA TGC CAG CAT CAT ACC TGC GGC |
| L138C-forward | CTG GGC CTG GGA CTG GGT GTG TGC CTG ATG CTG GGG CTG GAT TAT |
| L138C-reverse | ATA ATC CAG CCC CAG CAT CAG GCA CAC ACC CAG TCC CAG GCC CAG |
| A184C-forward | CAT AAT CTG CCG GAA GGT ATG TGC ATT GGT GTT AGC TTT GCA ACC |
| A184C-reverse | GGT TGC AAA GCT AAC ACC AAT GCA CAT ACC TTC CGG CAG ATT ATG |
| L200C-forward | GAT CTG CGT ATT GGT CTG CCG TGC ACC AGC GCC ATT GCA ATT CAG |
| L200C-reverse | CTG AAT TGC AAT GGC GCT GGT GCA CGG CAG ACC AAT ACG CAG ATC |
| A203C-forward | ATT GGT CTG CCG CTG ACC AGC TGC ATT GCA ATT CAG GAT GTT CCG |
| A203C-reverse | CGG AAC ATC CTG AAT TGC AAT GCA GCT GGT CAG CGG CAG ACC AAT |
| Q207C-forward | CTG ACC AGC GCC ATT GCA ATT TGC GAT GTT CCG GAA GGC CTG GCA |
| Q207C-reverse | TGC CAG GCC TTC CGG AAC ATC GCA AAT TGC AAT GGC GCT GGT CAG |
| A214C-forward | CAG GAT GTT CCG GAA GGC CTG TGC GTT GCC CTG GCA CTG CGT GCA |
| A214C-reverse | TGC ACG CAG TGC CAG GGC AAC GCA CAG GCC TTC CGG AAC ATC CTG |
| L217C-forward | CCG GAA GGC CTG GCA GTT GCC TGC GCA CTG CGT GCA GTG GGT CTG |
| L217C-reverse | CAG ACC CAC TGC ACG CAG TGC GCA GGC AAC TGC CAG GCC TTC CGG |
| V272C-forward | GCA GCG GGT GCA ATG ATT TTT TGC GTT AGC CAT GAA GTT ATC CCG |
| V272C-reverse | CGG GAT AAC TTC ATG GCT AAC GCA AAA AAT CAT TGC ACC CGC TGC |
| M295C- forward | ACC ACC GCA ACC GTT GGC CTG TGC GCA GGC TTT GCC CTG ATG ATG |
| M295C-reverse | CAT CAT CAG GGC AAA GCC TGC GCA CAG GCC AAC GGT TGC GGT GGT |
| **ZIP4 primers** | |
| A368C- forward | CTG CAG ACC TTC CTG AGC CTG TGC GTG GGT GCA CTC ACT GGG GAC |
| A368C- reverse | GTC CCC AGT GAG TGC ACC CAC GCA CAG GCT CAG GAA GGT CTG CAG |
| A368V- forward | CTG CAG ACC TTC CTG AGC CTG GTG GTG GGT GCA CTC ACT GGG GAC |
| A368V- reverse | GTC CCC AGT GAG TGC ACC CAC CAC CAG GCT CAG GAA GGT CTG CAG |
| A368F- forward | CTG CAG ACC TTC CTG AGC CTG TTT GTG GGT GCA CTC ACT GGG GAC |
| A368F- reverse | GTC CCC AGT GAG TGC ACC CAC AAA CAG GCT CAG GAA GGT CTG CAG |
| A514C- forward | CAC AAC TTC GCC GAC GGG CTG TGC GTG GGC GCC GCC TTC GCG TCC |
| A514C- reverse | GGA CGC GAA GGC GGC GCC CAC GCA CAG CCC GTC GGC GAA GTT GTG |
| A514V- forward | CAC AAC TTC GCC GAC GGG CTG GTG GTG GGC GCC GCC TTC GCG TCC |
| A514V- reverse | GGA CGC GAA GGC GGC GCC CAC CAC CAG CCC GTC GGC GAA GTT GTG |
| A514F- forward | CAC AAC TTC GCC GAC GGG CTG TTT GTG GGC GCC GCC TTC GCG TCC |
| A514F- reverse | GGA CGC GAA GGC GGC GCC CAC AAA CAG CCC GTC GGC GAA GTT GTG |
| A532C- forward | ACC GGG CTG GCC ACC TCG CTG TGC GTG TTC TGC CAC GAG TTG CCA |
| A532C- reverse | TGG CAA CTC GTG GCA GAA CAC GCA CAG CGA GGT GGC CAG CCC GGT |
| A532V- forward | ACC GGG CTG GCC ACC TCG CTG GTG GTG TTC TGC CAC GAG TTG CCA |
| A532V- reverse | TGG CAA CTC GTG GCA GAA CAC CAC CAG CGA GGT GGC CAG CCC GGT |
| A532F- forward | ACC GGG CTG GCC ACC TCG CTG TTT GTG TTC TGC CAC GAG TTG CCA |
| A532F- reverse | TGG CAA CTC GTG GCA GAA CAC AAA CAG CGA GGT GGC CAG CCC GGT |
| G543C- forward | CAC GAG TTG CCA CAC GAG CTG TGC GAC TTC GCC GCC TTG CTG CAC |
| G543C- reverse | GTG CAG CAA GGC GGC GAA GTC GCA CAG CTC GTG TGG CAA CTC GTG |
| G543V- forward | CAC GAG TTG CCA CAC GAG CTG GTG GAC TTC GCC GCC TTG CTG CAC |
| G543V- reverse | GTG CAG CAA GGC GGC GAA GTC CAC CAG CTC GTG TGG CAA CTC GTG |
| G543F- forward | CAC GAG TTG CCA CAC GAG CTG TTT GAC TTC GCC GCC TTG CTG CAC |
| G543F- reverse | GTG CAG CAA GGC GGC GAA GTC AAA CAG CTC GTG TGG CAA CTC GTG |
